# Top-down attention modulates auditory-evoked neural responses in neurotypical, but not ADHD, young adults

**DOI:** 10.1101/2021.02.11.430824

**Authors:** Jasmine A. C. Kwasa, Abigail L. Noyce, Laura M. Torres, Barbara G. Shinn-Cunningham

## Abstract

Individuals differ in their ability to selectively attend to goal-relevant auditory stimuli. People with Attention-Deficit/Hyperactivity Disorder (ADHD) in particular tend to show cognitive deficits associated with distractibility and inefficiencies in inhibition and attention. We hypothesized that people with ADHD would exhibit poorer performance and weaker neural signatures of attentional control when undertaking a challenging auditory task that required strong top-down attention. Neurotypical (N = 20) and ADHD (N = 25) young adults with normal hearing listened to one of three concurrent, spatially separated speech streams and reported the order of the syllables presented while we recorded electroencephalography (EEG). We tested both the ability to sustain attentional focus on a single “target” stream and the ability to monitor the target but flexibly switch attention to an unpredictable “interrupter” stream from another direction if and when it appeared. Although both stimulus structure and task demands affected behavioral performance, ADHD status did not. In both groups, the interrupter evoked larger neural responses when it was to be attended compared to when it was irrelevant, including for the P3a “reorienting” response previously described as involuntary. This attentional modulation was weaker in ADHD listeners, even though their behavioral performance was no lower. Across the entire cohort, individual performance correlated with the degree of top-down modulation of neural responses. These results demonstrate that listeners differ in their ability to modulate neural representations of sound based on task goals. Adults with ADHD have weaker volitional control of attentional processes than their neurotypical counterparts.

**Significance Statement:** ADHD and neurotypical listeners attended to one speech stream among distractors while neural responses were measured with electroencephalography. Behavioral performance varied with stimulus structure and task demands, but not with ADHD status. In both groups, top-down attention modulated stimulus-evoked neural responses: interrupting sounds elicited weaker responses when the sounds were ignored compared to when they were attended. This modulation affected a late “orienting” response (P3a) that has been previously described as automatic and not dependent on internal state. Importantly, ADHD subjects showed weaker attentional filtering than did neurotypical controls. At the individual level, performance correlated with neural metrics. Our results demonstrate that people vary widely in how flexibly they can use attention to modulate sensory responses based on task goals.

## Introduction

Competing sounds, like a teacher’s voice against the sudden trill of a cell phone, pose a challenge to the cognitive process of attention. Listening in such environments depends upon a push-and-pull between goal-directed attention, allocated to a source (the teacher’s voice), and automatic, involuntary shifts of attention to other salient, unexpected sounds (the ringing phone). The outcome of this attentional contest depends on the strength of an individual’s “top-down” control of attention relative to their susceptibility to “bottom-up” attentional capture (1, 2). In order to better understand cognitive control during auditory selective attention, we measured electrophysiological correlates of this push-and-pull dynamic by isolating measures of individuals’ top-down and bottom-up attention in a neurodiverse population.

Top-down attention enhances neural responses to attended stimuli and suppresses those to ignored stimuli (3, 4). Specifically, the magnitude of stimulus-elicited event-related potentials (ERPs) in electroencephalography (EEG) depends on attentional focus. The N1 response, a negative-going ERP component approximately 100 ms after stimulus onset, is larger when listeners focus top-down attention on the evoking source and smaller when they focus attention elsewhere (5–9). This difference yields a measure of top-down attentional control. Conversely, bottom-up attentional orienting to new events depends largely on their salience, rather than a listener’s goals. Salient events reliably elicit positive ERP responses approximately 300 ms after stimulus onset. This positive-going P3 response reflects attentional capture (10). The interplay between top-down focus and bottom-up salience ultimately determines what a listener truly “hears.” Importantly, individual differences in cognitive functioning affect the ability to focus on goal-relevant stimuli and orient appropriately to new stimuli (11, 12). These differences are reflected in large individual differences in the magnitudes of N1 and P3 responses to the same stimuli, even among neurotypical, normal-hearing adults.

People with Attention-Deficit/Hyperactivity Disorder (ADHD) struggle with tasks requiring top-down control, including selective attention (13–15). In laboratory environments, these individuals may perform just like age-matched controls; however, they often engage different, compensatory executive processes, accompanied by reduced neural activity in the regions recruited by neurotypical brains (16, 17). For example, children and adults with ADHD exhibit attenuated N1 responses; they also show attenuation of a subset of P3 responses related to decision-making (likely the P3b; see review in (18)). Thus, behavioral studies alone cannot identify important neural processing differences in ADHD, nor do they reveal whether selective attention deficits are due to poor top-down control of attention or to atypical bottom-up responses (19). To unravel the mechanistic roots of the disorder, one must characterize both behavior and brain activity, and measure neural responses that reflect both volitional, top-down attentional control and bottom-up, exogenous attention.

Here, we tested young adults with and without ADHD on a paradigm that stressed cognitive control of attention. We assessed both the ability to focus on a single stream of sound and the ability to flexibly switch attention from a target stream to a new, interrupting sound. We included key conditions in which the stimuli were identical between trials, but the task differed, altering only the internal state of the subjects. This allowed us to isolate effects of top-down attention on behavior. We used EEG to concurrently record neural responses and capture the N1 and the P3a, whose strengths predominantly reflect top-down and bottom-up attention processes, respectively. We reasoned that even if behavioral metrics did not differentiate ADHD from neurotypical subjects, the neural signatures of attentional focus might.

We found that ADHD subjects exhibited weaker top-down attentional modulation of neural responses to interrupting sounds than did neurotypical listeners, even though behavioral performance did not differ between groups. We also found that top-down attention modulates not only the N1, but the P3a, previously described as driven exclusively by bottom-up mechanisms (20). At the individual subject level, attentional modulation of both N1 and P3a responses correlated with behavioral performance. Together, these results demonstrate that individuals differ in their ability to control top-down attention in the face of salient interruptions, and that this ability is weaker in ADHD than in neurotypical subjects.

## Results

We recruited young adults (18–30 years old) with and without ADHD diagnoses to perform auditory selective attention tasks while we recorded concurrent EEG. On each trial, subjects began by focusing attention on a three-syllable “Target” stream of human speech, which was always diotic, with zero interaural time difference (ITD). The target consisted of the syllables /ba/, /da/, and /ga/, with the order randomly permuted from trial to trial (Figure 1). Every trial also contained a five-syllable “Distractor” stream (each syllable chosen randomly, with replacement, from the same set of syllables), which started after the Target and was spatialized to the right (ITD −700 *μ*s). Finally, two-thirds of trials contained a three-syllable “Interrupter” stream, which was a random permutation of /ba/, /da/, and /ga/, similar to the Target. The Interrupter was spatialized to the left (ITD 700 *μ*s) and either temporally overlapped with the Target (Early) or began after the Target syllables ended (Late).

**Figure 1.**
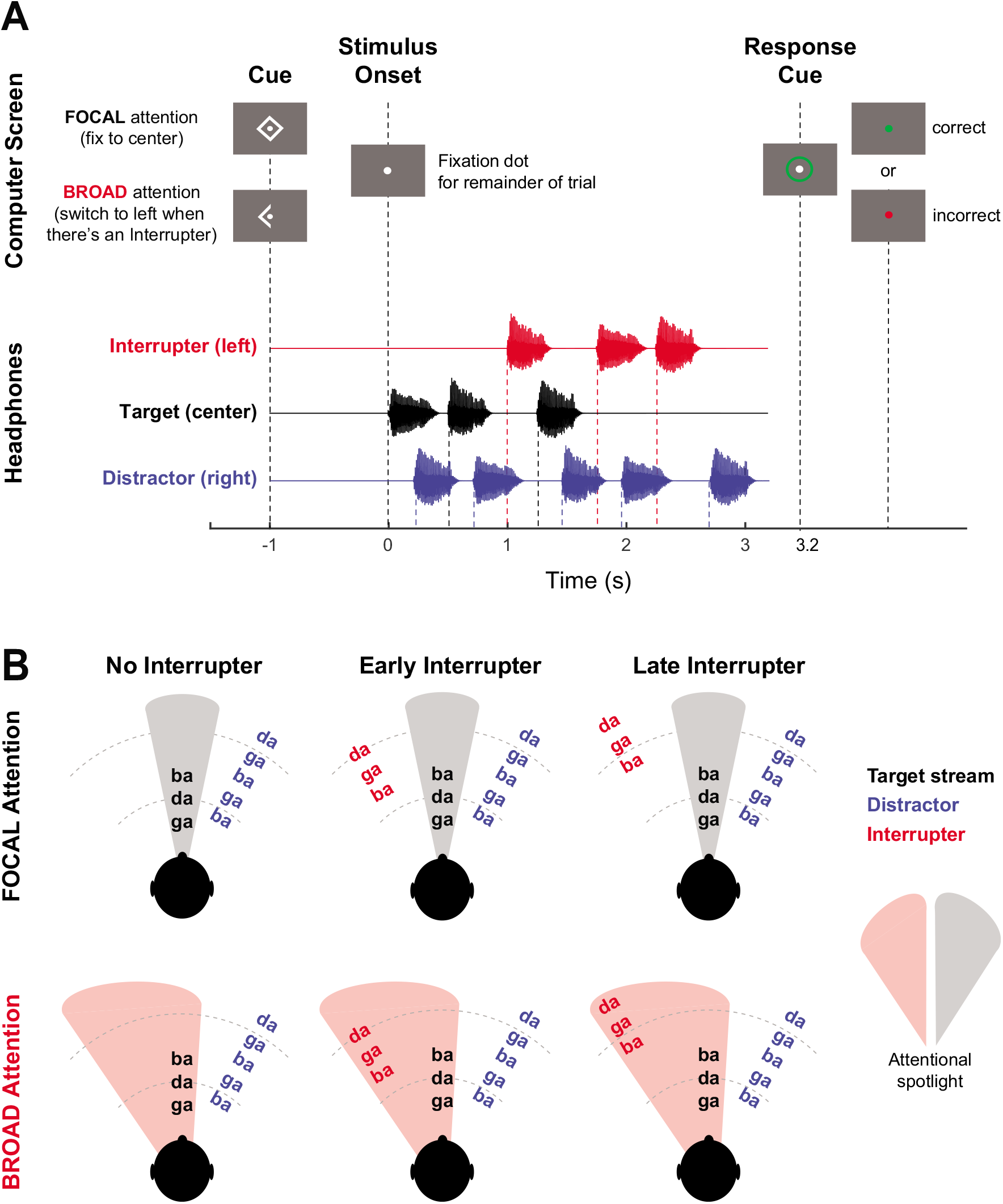
Experimental Setup. (A) Order of events in an “Early Interrupter” trial. A visual cue instructed subjects to either engage in FOCAL attention, monitoring only the central Target stream, or in BROAD attention, in which they needed to monitor for the onset of a left-lateralized Interrupter stream and, if an Interrupter occurred, switch their attention to it. (B) Schematic of our factorial experiment design. Across the columns are the approximate relative timing of the three stimulus streams in No Interrupter, Early Interrupter, and Late Interrupter conditions. In rows are the hypothesized attentional states required of the FOCAL and BROAD attention tasks.

On each trial, a visual cue instructed subjects to either ignore the Interrupter or to switch attention to it. On FOCAL trials, subjects were to maintain attention on the Target stream and, at the end of the trial, report the Target syllable order. On BROAD trials, subjects were to monitor the Target stream unless and until an Interrupter, coming from the left, occurred (Figure 1B). If an Interrupter occurred (2/3 of trials), subjects were to switch attention away from the Target and instead report the Interrupter’s syllables. On BROAD trials in which no Interrupter appeared (1/3 of trials), subjects were to simply maintain focus and report the Target syllables. BROAD attention trials, therefore, were particularly challenging, as subjects had to monitor the Target stream but be prepared to switch attention to the Interrupter stream if and when it played.

Because No Interrupter, Early Interrupter, and Late Interrupter trials were randomly intermingled, subjects could not anticipate whether or when an Interrupter would appear on a given trial, forcing them to adopt flexible listening strategies. Syllable timings in all streams were staggered so that event-related potentials evoked by many of the early syllable onsets could be temporally isolated, allowing us to analyze the modulatory effects of top-down attention on the neural representations of the corresponding syllables.

All behavioral analyses were performed on arcsine-transformed proportion-correct scores. All measures, both behavioral and neural, were analyzed using within-subject comparisons. See Materials and Methods for further details about the stimuli and analysis.

### Stimulus features and attentional focus, but not ADHD status, affect task performance

Subjects in both cohorts reported the correct syllable order at rates far above chance (mean = 62.3%, std. dev. = 15.3%; Figure 2). Accuracy was significantly higher in FOCAL than BROAD attention (F(1,41)= 85.4, p=1.42e-11). There was also a main effect of Interrupter Type (F(2,82) = 7.56, p=.00097), with lower accuracy on Late Interrupter trials than on either Early Interrupter (t(85) = 3.10, p=.008) or No Interrupter trials (t(85)=3.51,p=.002). This is likely due to the fact that the onsets of the Late Interrupter syllables aligned more closely in time with the onsets of syllables in the Distractor stream, resulting in greater perceptual interference (see Figure S1 for visualization of the syllable overlaps). Consistent with past studies, there was no main effect of ADHD status on performance [F(1,41)= 1.753, p = 0.193], and no significant interaction of ADHD status with other factors, or any other significant interactions.

**Figure 2.**
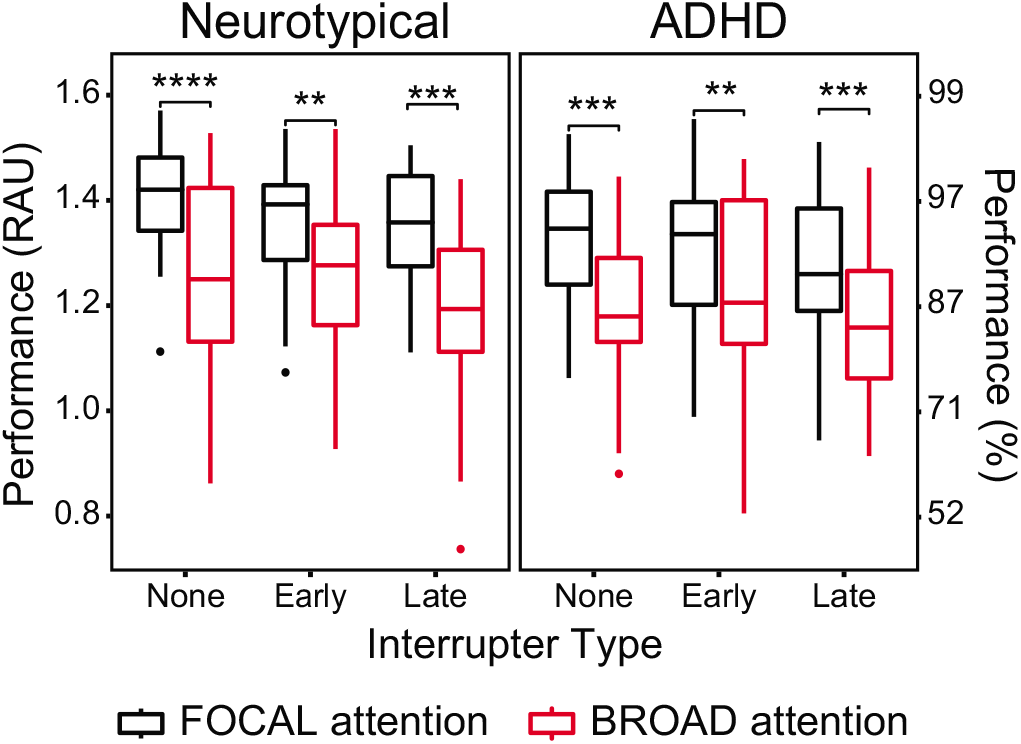
Performance for Neurotypical (left panel, N = 25) and ADHD (right panel, N = 20) groups in rational arcsine units for each Interrupter Type (None, Early, or Late), separately for FOCAL (black) and BROAD (red) attention conditions. Subjects performed worse in the BROAD condition compared to the FOCAL condition in all Interrupter Trial Types and for both ADHD and Neurotypical groups. This cost of broadening attention was present even in the No Interrupter trials, when there was no difference between FOCAL and BROAD in either of the stimulus features (or the correct response) and only the breadth of attention changed.

Performance was significantly higher on FOCAL than on BROAD attention trials at each level of Trial Type and ADHD status (post-hoc p<.001, Bonferroni adjusted for 6 comparisons). This is particularly interesting for the No Interrupter trials, in which subjects heard statistically identical stimuli and never had to shift attention away from the Target; these trials differed only in whether subjects were focusing exclusively on the target (FOCAL) or preparing to switch attention to an Interrupter (BROAD). Thus, there is a performance cost of broadening attention: subjects are more accurate in reporting the Target in FOCAL trials than in BROAD trials even when no Interrupter appears.

### Before an interrupter occurs, neural responses are similar for broad and focal attention and don’t differ across groups

We hypothesized that the amplitudes of neural responses might reflect the cost of broadening attention that we observed in accuracy. Specifically, we posited that Target-evoked N1 amplitudes might be smaller in BROAD trials compared to FOCAL because listeners who were anticipating a task-relevant Interrupter might be less focused on the Target. We analyzed neural responses to all Target syllables in No Interrupter and Late Interrupter trials (Figure 3A). (Early Interrupter trials were excluded because the Interrupter began while the Target was still ongoing.) N1 responses evoked by the Target syllables were not affected by Condition [F(1,41)= 0.264, p=0.610], ADHD status [F(1,41)= 0.028,p=0.867], or their interaction [F(1,41)= 0.155, p =0.695].

**Figure 3.**
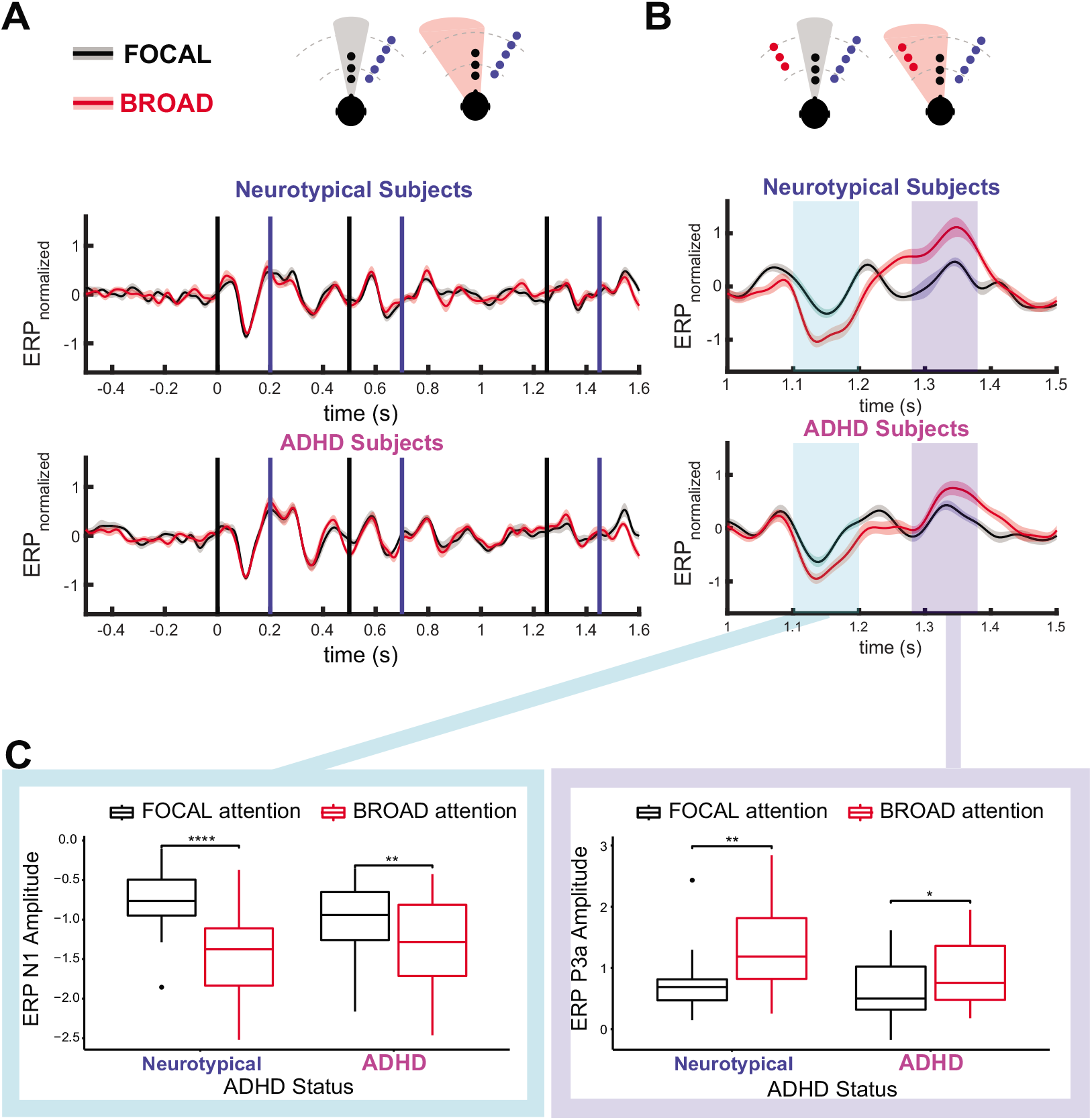
N1 and P3a responses. (A) Trial structures (top) and grand average ERP (bottom) for trials without an Interrupter in the first 1.5 seconds (i.e., No Interrupter and Late Interrupter trials). Event-related potentials are shown separated by ADHD status (top panel: Neurotypical, bottom panel: ADHD) and condition (FOCAL in black, BROAD in red, and error patches depict standard error). Vertical lines depict onset times for Target (black) and Distractor syllables (blue). (B) Trial structures (top) and grand average ERP (bottom) for the first Early Interrupter (t = 1.0 s). ERP peak amplitudes were calculated within the highlighted regions (left: N1; right: P3a). These traces are also shown separated by ADHD status (top panel: Neurotypical, bottom panel: ADHD) and condition (FOCAL in black, BROAD in red). (C) Boxplots showing ERP N1 (left) and P3a (right) amplitudes, separated by ADHD status (N = 25) and Condition (N = 20).

We also hypothesized that subjects with weaker top-down control of attention might be worse at filtering out responses to the always-ignored stream, the Distractor. To explore this, we analyzed N1s evoked by all Distractors in the No Interrupter trials. Although ADHD subjects exhibited slightly larger Distractor N1s than Neurotypical subjects, this difference was not statistically significant [F(1,41)= 1.979, p=0.167]. In addition, neither Condition [F(1,41)= 0.055, p=0.817] nor the interaction of ADHD Status and Condition [F(1,41)= 1.218, p=0.276)] significantly affected N1 amplitude. Individual differences in Target and Distractor N1 responses are shown in Figure S2.

Post-hoc, we performed a non-parametric cluster-based test (21) to find any other ERP components modulated by attention or ADHD status. We tested from 0 s, when the Target begins to play, to 1.5 s, before the Late Interrupter begins. We detected no time windows in which the FOCAL and BROAD attention conditions significantly differed in either the ADHD or Neurotypical groups (see traces in Figure 3A).

### ADHD subjects exhibit weak top-down modulation of neural responses to an interrupting event

We hypothesized that top-down attention would modulate N1s evoked by an Interrupter. We therefore analyzed the responses evoked by the first syllable in the Early Interrupter. (Subsequent onsets of the Early Interrupter and all onsets of the Late Interrupter could not be isolated from other temporally adjacent events; see Materials and Methods.) Overall, the first syllable of the Early Interrupter, which occurs before the final syllable of the Target stream, elicits larger N1s in BROAD than in FOCAL trials [F(1,41)=41.2, p<0.001; Figure 3B]. This demonstrates that volitional attention modulates this neural response, either by enhancing the Early Interrupter N1 during BROAD attention, suppressing it during FOCAL attention, or some combination of the two. Importantly, attention modulates this Early Interrupter N1 more weakly in ADHD subjects than in Neurotypical subjects. Statistically, this observation is supported by a significant Condition x ADHD Status interaction (F(1,41)=6.79, p = 0.0130), even though there was no main effect of ADHD Status [F(1,41)=0.002, p=0.968]. This result demonstrates that ADHD subjects modulate the neural responses evoked by the Interrupter based on task demands less than do neurotypical listeners, consistent with weaker top-down control of attention.

### Attentional focus modulates the P3a, usually considered a bottom-up response

Prior work suggests that the P3a ERP component is elicited by unexpected stimuli, exogenously, and is influenced only by stimulus features (20) — although at least one study shows that other cognitive disorders affect P3a responses (22). We thus hypothesized that this bottom-up response would be stronger in ADHD subjects due to their high distractibility (23) and did not expect attentional state to affect the magnitude of the response in either subject group. Unexpectedly, we found that the P3a elicited by the first onset of the Early Interrupter was modulated by attention in both ADHD and Neurotypical subjects. Specifically, the Early Interrupter P3a was larger in the BROAD condition, when the Interrupter was behaviorally relevant, than in the FOCAL condition, where it was to be ignored [F(1,42)=19.8, p=6.1e-05]. Rather than finding overall larger raw P3a amplitudes in ADHD subjects due to weaker attentional filtering, we found no main effect of ADHD status [F(1,42)= 2.52, p=0.12)]. While top-down modulation of the P3a tended to be weaker in ADHD than Neurotypical subjects, similar to the N1 response modulations, the difference did not reach statistical significance [Condition x ADHD Status interaction: F(1,42)=3.62, p=.064; see Figure 3C].

### At the individual level, the degree to which neural responses change with task goals correlates with behavioral performance on the task

While we find significant group differences in the strength of top-down modulation of the N1 evoked by the first syllable of the Early Interrupter, there is also significant individual variability of these responses within each group and significant overlap across groups in both behavioral and neural measures. We therefore tested whether there is a relationship between task performance and attentional modulation of the Early Interrupter N1 and the P3a at the individual subject level. For each subject, we calculated the degree of attentional modulation for the Interrupter N1 (ΔN1 = N1 peak in FOCAL - N1 peak in BROAD) and P3a (ΔP3a = P3a peak in BROAD - P3a peak in FOCAL), each defined so that large positive values indicate strong changes in neural responses due to top-down attention. Both ΔN1 and ΔP3a significantly and positively correlate to accuracy on the Early Interrupter trials (r(41) = 0.315, p = 0.040 and r(42) = 0.560, p = 7.81 x 10^−5^, respectively; Figure 4), demonstrating that individual differences in top-down control of attention relate directly to differences in performance.

**Figure 4.**
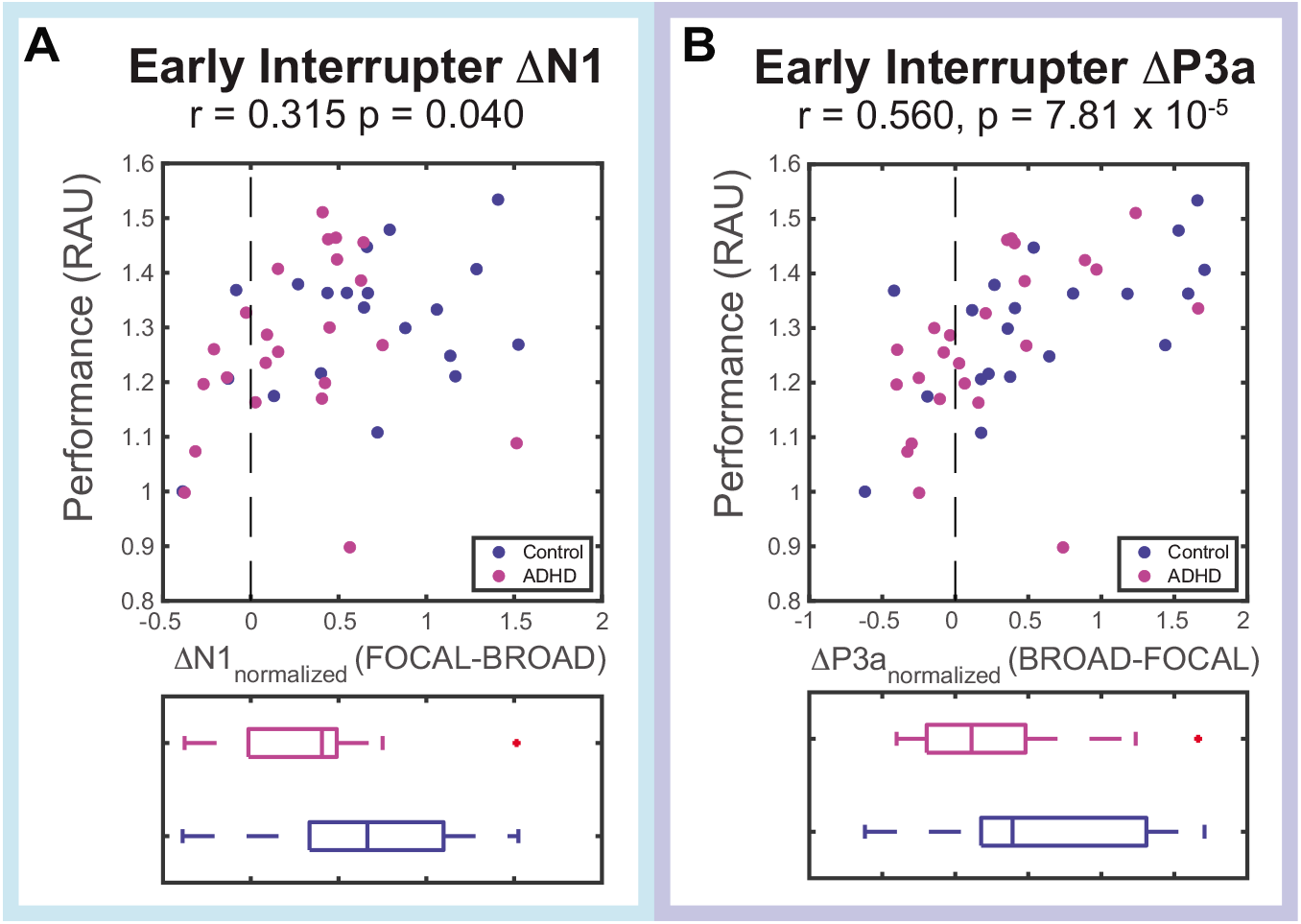
(A) Individual differences in total Early Interrupter trial performance plotted against individuals’ ΔN1s (FOCAL-BROAD) to the first Early Interrupter, depicting a significant correlation between the attention modulation and performance. The bottom panel shows the spread of individuals’ ΔN1 according to the population (ADHD in magenta and Neurotypical in blue). (B) Same data as (A) but for ΔP3a (BROAD-FOCAL). The correlation between attention modulation and performance is strongly significant. ADHD status significantly affects ΔN1(t(41)=2.61, p = 0.0127) and marginally affects ΔP3a (t(42)=1.90, p=0.0640).

## Discussion

Young adults with ADHD typically struggle with inattention, impulse control, and hyperactivity. We hypothesized that this population’s atypical cognitive strategies would yield poor auditory selective attention performance, especially in cases where unpredictable, salient interrupting events drew attention. We designed a demanding auditory experiment that exercised different attentional demands to explore how top-down attention control interacted with bottom-up attention capture in neurotypical and ADHD subjects. Although we did not observe group differences in task performance, we did find differences in how strongly attentional focus modulated neural responses, with ADHD subjects exhibiting weaker modulation. This pattern aligns with past work describing compromised preparatory-related neural responses in ADHD and reduced activation of networks involved in both bottom-up and top-down processing, even when there are no behavioral group differences (16, 17, 19).

We did not observe any attentional modulation, in either subject group, of the neural responses elicited in the early portion of trials. This likely reflects the similarity of task demands in this period. In all conditions, listeners had to initially focus on the Target and ignore the Distractor; only towards the end of a trial, if an Interrupter appeared, did they sometimes have to switch attention from the Target. This likely explains why Distractor- and Target-evoked ERPs early in the trial are similar for FOCAL and BROAD conditions.

The N1 response’s peak magnitude is strongly modulated by top-down attention (8, 24–26), and the degree of this modulation correlates with performance on selective attention tasks (27). We hypothesized that N1 modulation would be reduced in ADHD subjects, reflecting reduced efficacy of attentional control. We contrasted N1 responses elicited by FOCAL trials, where listeners always reported the Target stream, and BROAD trials, where they began listening to the Target but had to be prepared to switch attention to an Interrupter, if it occurred. We found strong attentional effects on the magnitude of N1 responses evoked by the salient and unpredictable Early Interrupter. The N1 was larger in BROAD trials, presumably because listeners needed to shift focus to the Interrupter, than in FOCAL trials, where the Interrupter was not task relevant. This difference was reduced in subjects with ADHD, suggesting that ADHD listeners are less able to use top-down attention to modulate neural responses to salient, interrupting events. This supports the account of ADHD as a deficit of or alternative solution to executive functioning.

We found that for both neurotypical and ADHD subjects, the P3a evoked by the Early Interrupter was stronger in the BROAD than the FOCAL trials. This was unexpected. Past studies discuss the P3a as a response evoked by salient events, such as a change to a repeated stimulus, or the occurrence of a novel, task-irrelevant sound (8, 28–30). P3a amplitudes can vary with age and circadian arousal levels (31), the frequency and type of the target stimulus in relation to ignored stimuli, personality (32), cognitive maturation (33), and fatigue due to time spent on task (34). Some studies report that the amplitude of P3a responses elicited by a task-irrelevant sound is attenuated by increased cognitive load in a primary task (32, 33 but see also 34, 35). However, this cannot account for our results. In our experiment, we observe smaller P3a responses in the FOCAL condition. If anything, this condition places less cognitive load on our listeners than the BROAD condition, which requires listeners to be ready to reorient attention to the Interrupter; a cognitive-load account would predict that the P3a evoked by the Interrupter should be greater in the FOCAL condition than the BROAD condition. Instead, our results suggest that when listeners focus attention on the Target and suppress distracting events, it leads to a suppression of both the N1 and the P3a evoked by a to-be-ignored stimulus.

In the current study, ADHD subjects trend toward weaker attentional modulation of the P3a compared to neurotypical listeners. Some past studies have reported relationships between raw P3a amplitude and clinical measures of cognitive ability (39–41). However, we are not aware of any previous studies examining the efficacy of top-down attention in modulating the P3a. Our finding further supports the idea that ADHD manifests as a reduced ability to deploy top-down attentional control. Future studies should directly address this ability in this population.

Finally, the group differences we found are significant, despite large individual subject differences. These individual differences are not random; instead, performance correlates with how strongly top-down attention modulates the N1 and P3a evoked by interrupting events. Listeners who perform best are those who more strongly suppress neural responses to an event when it is task irrelevant compared to when it is task relevant. This correlation suggests that the differences in the strength of top-down attentional control relate directly to differences in behavioral ability and thus may provide a more nuanced measure of the ability to ignore salient distractions than does a categorical label.

Scientifically, our results suggest at least two issues that deserve additional investigation. First, we need to further explore the influence of top-down attention on the P3a response to confirm such effects, for instance, by isolating the response in different experimental paradigms. Second, further research into the mechanisms that lead to individual differences in top-down control should be conducted. One avenue for future research is to explore whether there are signatures of ADHD in oscillatory brain activity. For instance, the ratio between theta waves (4-8 Hz) and beta waves (12-20 Hz) has shown promise as a neural marker of ADHD, but not one sensitive or selective enough to be clinical relevant (42–46). Similarly, lateralization of parietal alpha (8-14 Hz) oscillations are associated with spatially directed attention, with increases in alpha power in the hemisphere representing to-be-ignored visual or auditory events (47–50).

### Clinical implications for ADHD and caveats

Our results support the idea that behavioral assessments are less sensitive than neural measures of ADHD (23, 51, 52). Still, as in many prior studies, our effect size is too small to be clinically useful for diagnosis or prognosis (42–45).

Increased distractibility in the ADHD population has been well-documented in both research and clinical settings (23), as has disrupted preparatory processing and performance monitoring in psychophysical tasks. The under-arousal theory of ADHD proposes that these issues are due to compromised attentional orienting in these individuals (53, 54). This account is consistent with our finding that ADHD listeners are less able to modulate neural responses evoked by salient, task-irrelevant interrupters than are neurotypical subjects. We argue that, compared to controls, ADHD subjects are less effective at filtering out salient but irrelevant events; ADHD biases the attentional systems to focus on salient stimuli, regardless of their behavioral importance.

It is worth noting that we did not separate our ADHD sample into the three subtypes of ADHD (inattentive, hyperactive/impulsive, and combined) identified in the DSM-5. Executive function deficits greatly vary within the ADHD population (55, 56), and this could explain some of the inter-subject variability amongst ADHD subjects. Similarly, ADHD status in our sample was determined by self-report of previous diagnosis. The heterogeneity of the sample was exacerbated since subjects had been diagnosed by different physicians and psychologists at different developmental points, which in turn likely affected coping mechanisms and comorbidities. Finally, our sample was largely comprised of students from relatively high socioeconomic backgrounds and enrolled in a highly selective American university. Future work should consider these factors and attempt to understand their role in ADHD and attention.

### Conclusions

During a demanding auditory task, neural responses evoked by an unpredictable interrupting sound are larger when that interrupter is behaviorally relevant than when it is to be ignored. This modulation affects not only the sensory N1 response, but also the P3a, which has been previously thought of as an automatic orienting response. Individual differences in how well listeners perform on this demanding auditory task correlate with how strongly they modulate neural representations based on task demands. Despite substantial overlap between neural signatures of attentional control in ADHD and neurotypical listeners, group differences are significant: ADHD listeners demonstrate weaker top-down modulation of neural responses evoked by an unpredictable interrupting sound. Future work should be undertaken to replicate the effects of top-down attention on the P3a, and to explore whether the group differences we see between ADHD and neurotypical listeners manifest similarly in attentionally demanding tasks in other sensory modalities.

## Materials and Methods

The Boston University Internal Review Board approved all study procedures. All participants gave written informed consent.

### Participants

We recruited 95 volunteers to complete an online intake survey (via Qualtrics, Provo, UT) reporting demographics, mental health and drug use history, and current mood and anxiety (Figure 5). Subjects who met inclusion criteria (ages 18-30 years, normal or corrected vision, and normal hearing) were invited to complete a lab visit. They gave written informed consent, completed a hearing screening, and performed an abbreviated practice test of the auditory experimental task.

**Figure 5.**
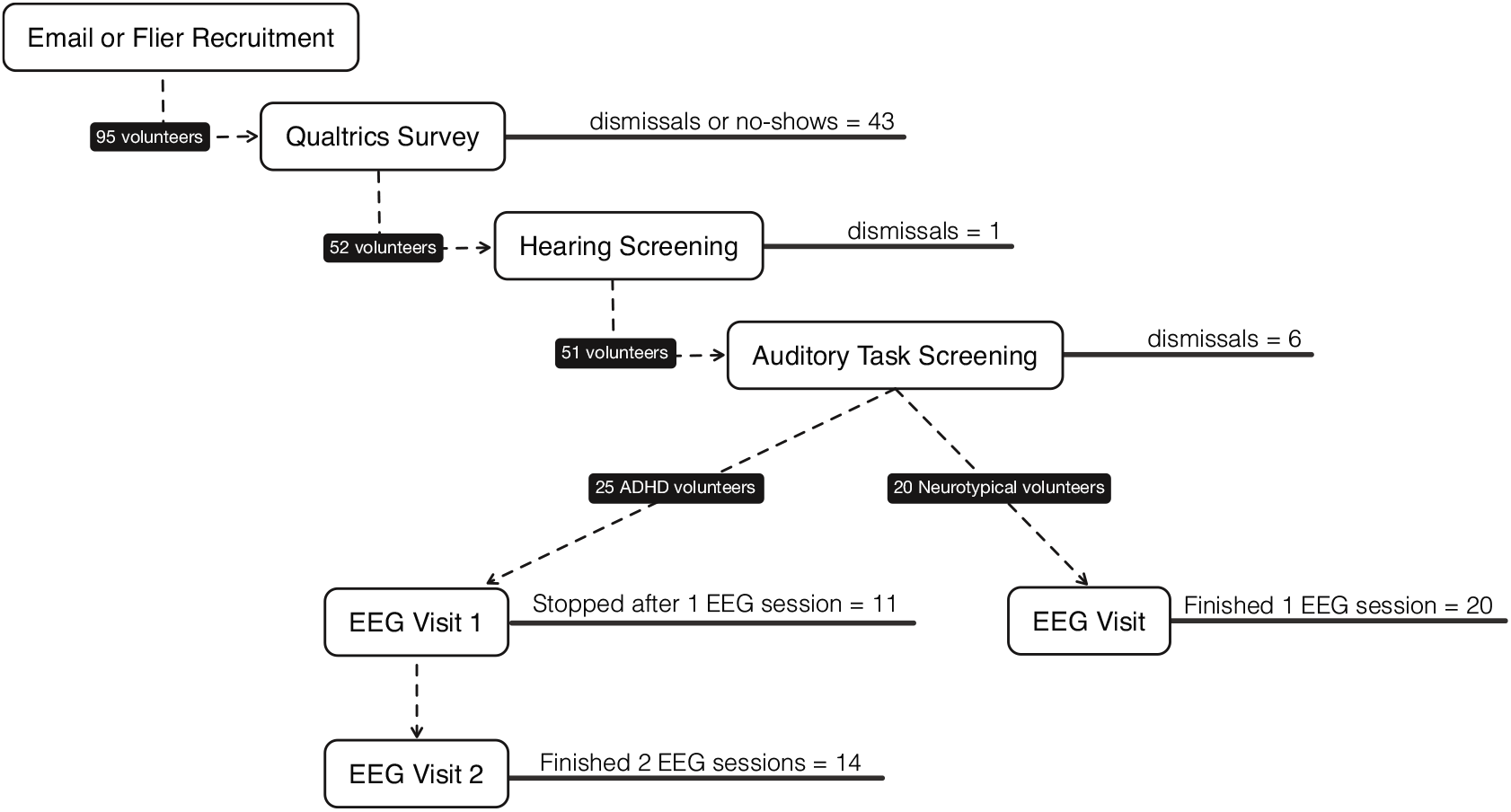
Subject numbers for all phases of the study. In this paper, we present data from 45 subjects, 25 with ADHD and 20 without a prior diagnosis, who are labelled Neurotypical.

Criteria for continuing in the study were auditory detection threshold levels at or below 20 dB HL for pure tones between 250 and 8,000 Hz (in octaves) and performance at or above 66% correct on the practice test. Figure 5 shows the recruitment pipeline and subjects retained or lost to follow-up at each stage. Of the 45 subjects who continued to the main experiment and had useable data, 25 self-reported that they had been diagnosed with ADHD. We refer to these participants as ADHD subjects (19 female, 6 male; age 21.4 +/- 2.7) the remaining 20 we call Neurotypical subjects (14 female, 6 male; age 21.8, 3.0). All subjects were compensated for their time and offered bonuses for good task performance.

ADHD subjects were initially tested while either on or off their stimulant medications, assigned randomly. A subset of the ADHD subjects (N=14) completed a second session of the same experiment on a subsequent day in the other medication state. Nine of these subjects performed on-medication first; five performed off-medication first. Post-hoc, we identified a strong practice effect: performance almost always increased on the second day, regardless of medication status. Therefore, in the main text we focus on data from each subject’s Day 1 data. The Supplemental Information shows results for all ADHD subject data from both days (Figures S1-3), highlighting within-subject comparisons (on vs. off medication) for the 14 participants who completed both experimental sessions.

### Auditory Experiment

#### Stimuli

Stimuli were streams of human speech comprising sequences of /ba/, /da/, and /ga/ syllables. Syllables were recorded by a male, native English speaker using an AudioTechnica AT4033 large diaphragm condenser microphone (Audio-Technica U.S. Inc., Stow, OH) in a sound-treated booth. These plosive syllables were selected because their abrupt onsets elicit strong ERPs. Individual syllables were recorded in isolation, cropped to be 437 ms long, and then concatenated with inter-stimulus intervals (ISIs) to form streams of randomly permuted syllables. Sound stimuli were presented via Etymotic ER-1 insert headphones (Etymotic, Elk Grove Village, IL). Syllables were spatialized to one of three stream locations using interaural time differences (ITDs) of 700 *μ*s (left of center), 0 *μ*s (center), or −700 *μ*s (right of center). Stimulus creation and experimental control were via custom software created in MATLAB (The MathWorks Inc, Natick, MA) using the PsychToolbox (57, 58).

#### Experiment design and task

Every trial contained a three-syllable “Target” stream heard from the center and a five-syllable “Distractor” stream spatialized to the right. The Target always began playing first, followed 200 ms later by the Distractor (Figure 2A). The Target syllable onsets were always presented at 0, 0.5s, and 1.25s. The Distractor syllable onsets were presented with one of two “rhythms:” 0.2s, 0.7s, 1.45s, 1.95s, and 2.7s (rhythm 1) or 0.2s, 0.95s, 1.45s, 2.2s, and 2.7s (rhythm 2). Two-thirds of trials contained a third “Interrupter” stream which was spatialized to the left and began either 1 s (Early Interrupter) or 1.5 s (Late Interrupter) after the Target. We balanced the design so there were equal numbers of No Interrupter, Early Interrupter, and Late Interrupter trials (Figure 2B). Each Interrupter was created to have one of two syllable rhythms, with onset times of 0s, 0.5s, 1.25s (rhythm 1) or 0s, 0.75s, 1.25s (rhythm 2). These were then delayed overall by either 1s, to create an Early Interrupter, or 1.5s, to create a Late Interrupter, before being added to the Target and Distractor. In all trials with Interrupters, only Distractor rhythm 1 was used to reduce the amount of overlap between competing syllables. All syllable timings are depicted in Figure S4).

Subjects were instructed to keep their eyes open and focused on a central fixation dot. Each trial began with a visual cue indicating the attentional state required. On FOCAL attention trials (diamond cue), subjects were to maintain attention on the Target and report the order of the /ba/, /da/, and /ga/ syllables presented in it. On BROAD attention trials (left-pointing arrow cue), subjects were to attend to the Target unless and until a (left-lateralized) Interrupter occurred. If an Interrupter began, subjects were to reorient attention to it and report its syllables, in order. The BROAD attention condition, therefore, was particularly challenging, as subjects had to both monitor the Target and be prepared to switch their attention to the Interrupter if it appeared (which could happen either Early or Late in the trial). Subjects were to always ignore the (right-lateralized) Distractor. Note that 1/3 of both FOCAL and BROAD attention trials were No Interrupter trials, letting us test the effect of attentional state in the absence of an attentional shift.

After all stimuli ended (3.2 s after the Target onset), a circle appeared at the fixation point to cue subjects to respond. Listeners withheld responses until this signal to prevent motor planning and motor artifacts from distorting the sensory-evoked EEG responses. We did not analyze reaction times.

Subjects reported back the required sequence (either Target or Interrupter, depending on the trial) using the keys 1, 2, and 3 to indicate the order of /ba/, /da/, and /ga/ syllables, respectively. Each subject trained on this mapping before the experiment began. After entering their response and before the start of the next trial, subjects received feedback as to whether or not their response was correct. In addition to hourly pay, subjects were given a $0.01 bonus for each correct response during the experiment (maximum bonus: $4.80).

### Behavioral performance analyses

We calculated the proportion of trials that were correctly reported. A trial was labeled correct if the subject correctly identified, in order, each of the three syllables in the proper stream (either the Target stream or Interrupter stream, depending on the trial) (59).

We used R (R Development Core Team 2011) and the rstatix, tidyverse, and ggpubr packages to perform mixed within/between 3-way ANOVAs to assess the effects of ADHD status, attention condition (FOCAL vs BROAD), and Interrupter Type (Early, Late, or None) on behavioral performance on the task. There were no extreme outliers, the data were normally distributed (Shapiro-Wilk p > 0.05), and there was adequate homogeneity of variances (Levene p > 0.05).

### EEG Acquisition

Subjects performed the experiment in front of an LCD monitor in a sound-treated booth. A BioSemi ActiveTwo system and accompanying ActiveView acquisition software recorded EEG from 64 channels arranged in the standard international 10-20 setup. Data were sampled at 2048 Hz. Auditory stimuli were presented through Tucker-Davis Technologies System 3 (TDT, Alachua, FL) hardware, which also inserted time-locked event flags into the EEG recording. Three external electrodes collected EOG responses from eye movements: two beside the eyes and one below the left eye.

### EEG Processing

Continuous EEG data were referenced against the average of the mastoid channels, then downsampled to 256 Hz. Data were bandpass filtered between 1 and 20 Hz with a zero-phase Kaiser filter to remove slow drift below 1 Hz and high-frequency noise above 20 Hz, including line noise. Independent components analysis (EEGLAB, 57) allowed us to isolate eye blinks, saccades, and other artifacts; components corresponding to such artifacts were identified by inspection and projected out of the data. Data from each trial were then epoched from one second before the visual cue onset to the end of the presentation period (4.5 s). Any processed epochs with amplitudes exceeding ±100 μV were rejected from further processing. Datasets with 3 or fewer non-adjacent, erratic channels (determined by visual inspection of ICA topographies and raw signal traces) were kept, and data from the bad channels interpolated (60). A final visual inspection removed any remaining contaminated trials.

### Event Related Potential (ERP) Calculations

The ERP components that we report are the peak magnitudes of the N1 and P3a responses. For each subject, trial type, and condition, we computed an average ERP across a broad cluster of 10 fronto-central channels, where auditory-evoked responses tend to be maximal (Fz, FC1, FCz, FC2, C1, Cz, C2, CP1, CPz, and CP2). Individuals’ N1 peaks were calculated by averaging epochs of EEG from trials of the same type and condition, then employing a custom peak-finding algorithm to identify the peak negativity in a window from 75 to 150 ms after each stimulus onset in all the streams. Individuals’ ERP P3as were calculated similarly, but for a peak positivity in a window from 280 ms to 380 ms after stimulus onsets. The full experiment comprised 240 trials, leaving each condition and trial type combination with a maximum of 40 trials for ERP calculation before artifact rejection. Because the number of trials that remained after artifact rejection varied across subjects and conditions, we used a Monte Carlo downsampling procedure to obtain usable ERP component estimates. For each subject, we calculated that subject’s minimum number of valid trials across all conditions (mean = 26.2, std dev = 5.50), then randomly selected this number of trials per condition for that subject. We repeated this procedure 100 times for each subject and then assigned the median N1 and P3a values calculated over all samples to that particular subject, condition, and trial type.

### ERP Statistical Analyses

ERP peaks were used in four hypothesis-driven analyses. To test the effects of ADHD Status and Attention Condition on task-relevant Targets and task-irrelevant Distractors, we included all Target-elicited N1s in No Interrupter and Late Interrupter trials, and all N1s elicited by Distractor onsets in No-Interrupter trials. To test the effects of ADHD Status and Attention Condition on the process of shifting attention to the Interrupter, we computed N1 and P3a peaks elicited by the first Interrupter onsets in Early Interrupter trials. Other onsets were contaminated by temporally adjacent stimuli, precluding extraction of the corresponding ERP components.

For each analysis, we computed each subject’s mean peak amplitude in each condition, and again used mixed within/between ANOVA to statistically test for differences.

We followed up with a non-parametric permutation test to further identify significant differences between the two Attention Conditions in No Interrupter and Late Interrupter trials up until t = 1.5 (21). For each subject, a paired sample t-value was calculated for each time point between the trial-length ERP for each Attention Condition. A null distribution for the t-test was derived for each subject from 1000 bootstrapped permutations of their data in which we swapped time points between the Attention Conditions. Time clusters over a certain pre-defined threshold were labeled significant.

## Acknowledgments

This research was funded by the National Institute on Deafness and Other Communication Disorders under Award Number R01 DC013825, as were ALN and BGSC. JAK was funded by the National Institute of Neurological Disorders and Stroke under Award Number F99 NS115331, National Science Foundation Graduate Research Fellowship Program under Grant Number DGE-1247312, and a Ford Foundation Pre-doctoral Fellowship. We would also like to thank the Boston University Center for Anxiety and Related Disorders, particularly Dr. Bonnie Wong and Emma Weizenbaum, for their expert advice and time.

