## Supplemental Information for "Top-down attention modulates auditory-evoked neural responses in neurotypical, but not ADHD, young adults"

**This PDF file includes:**

Supplementary text
Figures S1 to S4

### Supplementary Information Text

#### Medication status and day effects

We initially sought to explore the effects of stimulant medication on ADHD subjects' attentional functioning in addition to the factors explored in the main text, Condition and Interrupter Type.

All ADHD subjects who participated had been previously prescribed stimulant medications (e.g., methylphenidate, amphetamine salts, etc.) and provided a medication bottle as proxy for an ADHD diagnosis and to confirm that the medication was legally prescribed to that individual. Due to attrition between visits, only a small subset of ADHD (N=12) performed the task on two separate days once while on their prescribed stimulant medication (ADHD+) and once while abstaining (ADHD-), assigned randomly. See Figure S2 for a comparison of performance in these paired data.

A repeated measures ANOVA with Experimental Day, Medication Status, Interrupter Type, and Attention Condition as factors showed that only Day and Condition were significant (Day: $F(1) = 4.56$, $p = 0.0344$; Condition: $F(1) = 27.2$, $p = 6.61 \times 10^{-7}$; Interrupter: $F(2) = 2.50$, $p = 0.0856$; Medication: $F(1) = 3.44$, $p = 0.0657$). Specifically, performance improved significantly on the second day, regardless of medication use and the FOCAL condition was easier than the BROAD (as shown in the main text, Figure 2). Since medication status did not significantly improve performance, all analyses shown in the main text are derived from all subjects only on Day 1--Controls, ADHD+, and ADHD.

This removes any influence of training and groups all ADHD subjects, regardless of their Day 1 medication. These adaptations simplified our models to only include Condition and Trial Type. Re-running all analyses as ANOVAs without Day and Medication status, Condition retained its significance in Performance, N1 measures, and P3a measures.

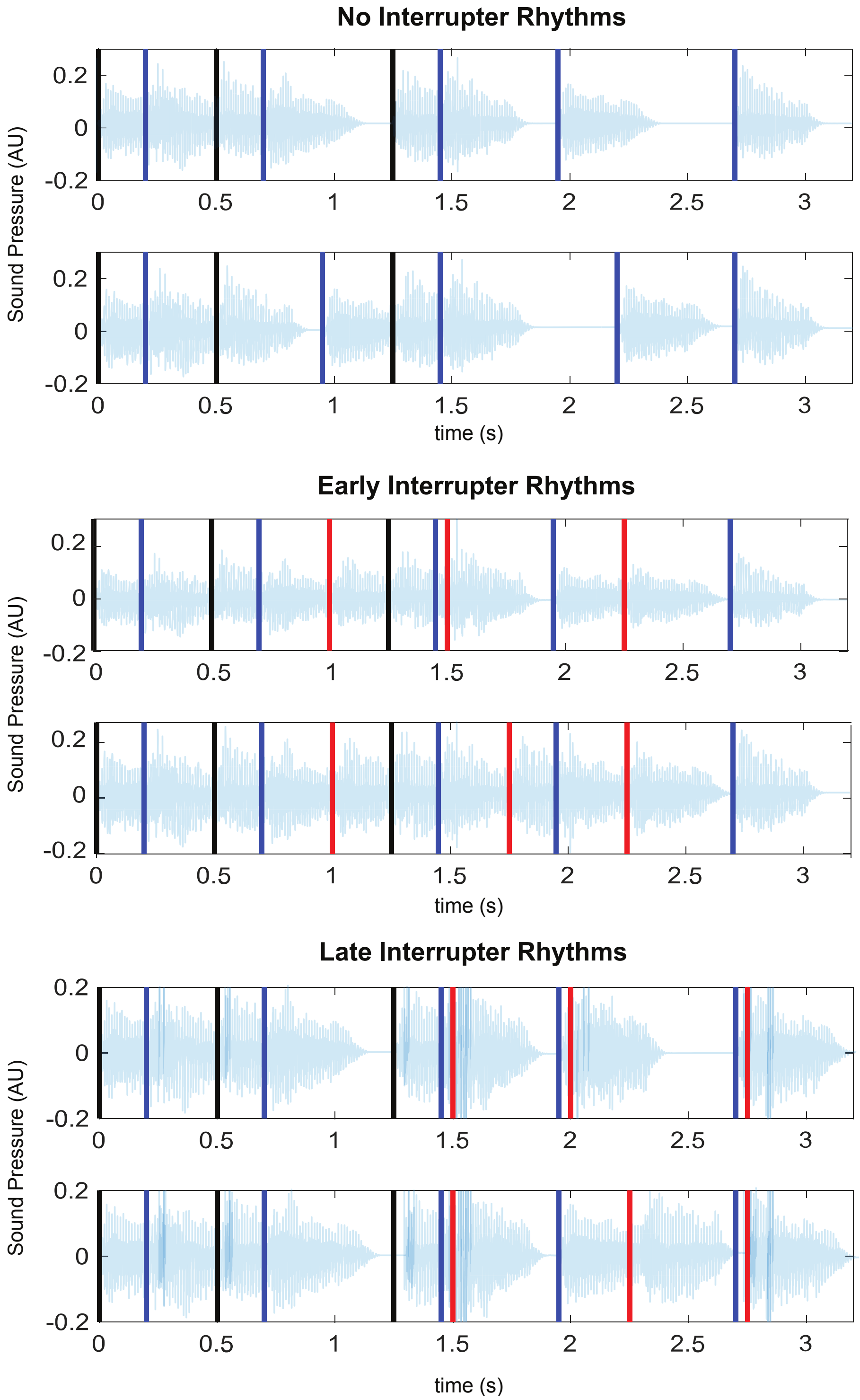


**Fig. S1** Stimulus timings for each Trial type, including the two possible "rhythms" per trial type used to encourage subjects to rely on attention to focus on Target or Interrupter stimuli. Sound pressure waves are depicted in light blue traces, while all onset timings are depicted as vertical lines: Target in black, Distractors in blue, and Interrupters in Red. The Late Interrupters were close in timing to Distractors, which prevented us from analyzing their corresponding ERP responses.

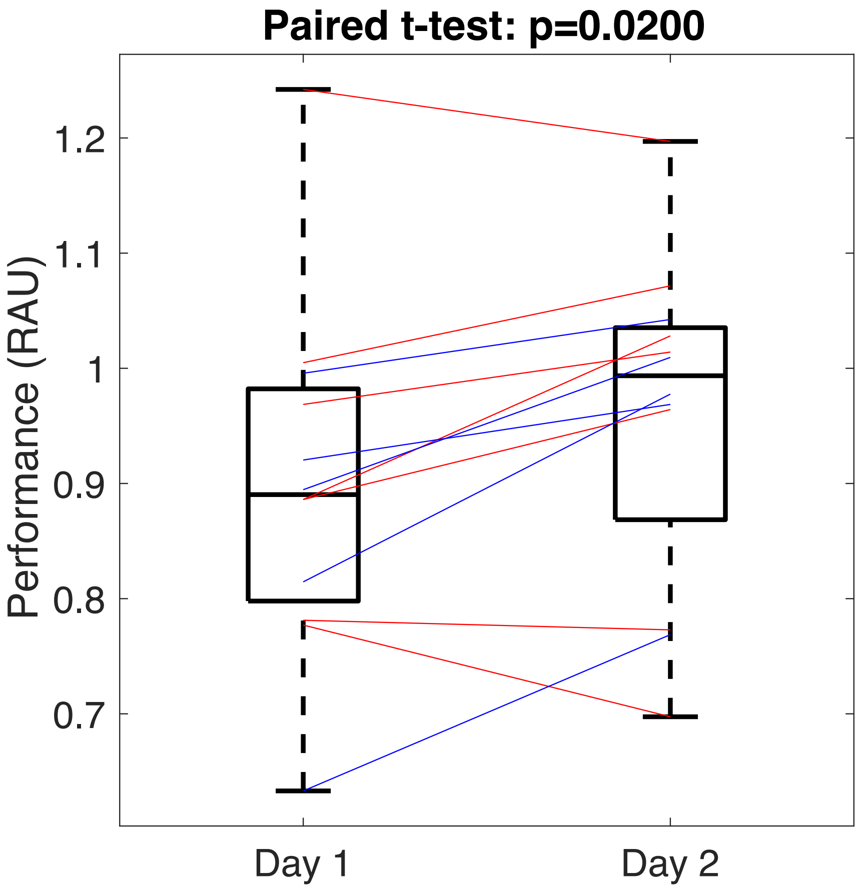


**Fig. S2.** Training and medication effects on behavioral performance. Twelve ADHD subjects completed the attention task on two separate days. Performance accuracy in rational arcsin units (RAU) is depicted in boxplots, with lines connecting performance of individual subjects. Blue lines are subjects who took their medications on Day 1 and red lines indicate medication on Day 2. A two-way ANOVA shows that the day effects (interpreted as training effects) are stronger than medication affects: performance significantly increases on Day 2 regardless of medication usage.

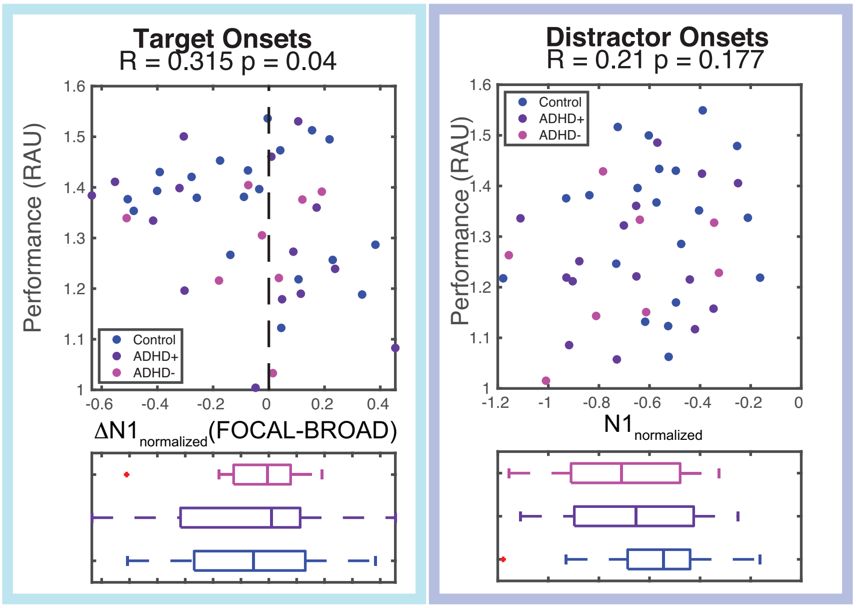


**Fig. S3.** (A) Individual differences in total task performance plotted against individuals' ΔN1s (FOCAL-BROAD) to Target onsets from trials in which an Interrupter does not interfere (No Interrupter and Late Interrupter trials). There is no correlation between attention modulation and performance. The bottom panel shows the spread of individuals' ΔN1 according to the population: ADHD participants abstaining from stimulant medication in magenta (ADHD-), ADHD participants using medication in purple (ADHD+), and neurotypical Controls in blue. (B) Individual differences in total task performance plotted against individuals' N1s evoked by all five Distractor onsets from trials in which no Interrupter appears (No Interrupter trials only). No correlation was found between Distractor N1 amplitude and performance.

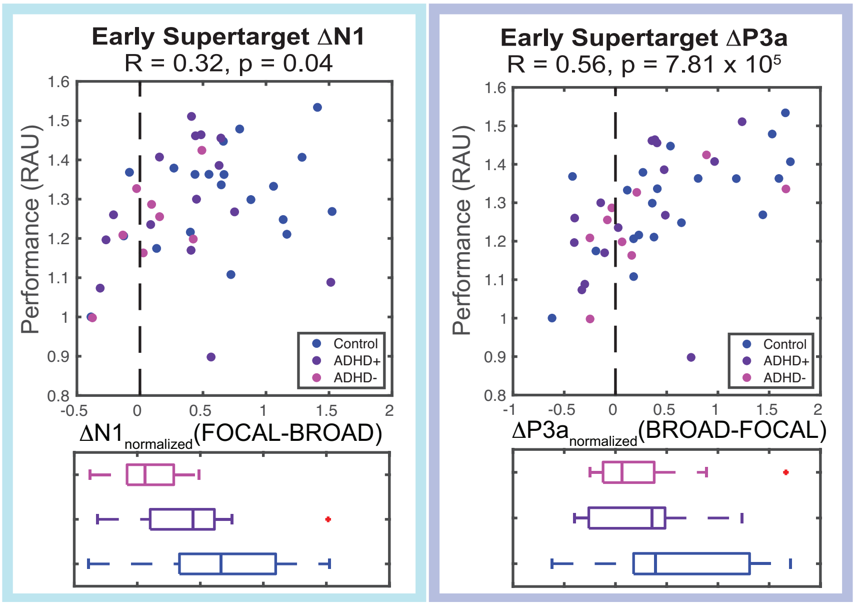


**Fig S4.** Individual differences in total Early Interrupter trial performance plotted against individuals' ΔN1s (FOCAL-BROAD) for all subjects: ADHD participants abstaining from stimulant medication in magenta (ADHD-), ADHD participants using medication in purple (ADHD+), and neurotypical Controls in blue. Lines connect data from unique individuals who took testing twice (N=12). Similar to the results of the same analysis with only Day 1 data (Figure 4), there is no correlation between attention modulation and performance. The bottom panel shows the spread of individuals' ΔN1 according to the population. (B) Individual differences in total task performance plotted against individuals' N1s evoked by all five Distractor onsets from trials in which no Interrupter appears (No Interrupter trials only). No correlation was found between Distractor N1 amplitude and performance.
